# RNF219 regulates CCR4-NOT function in mRNA translation and deadenylation

**DOI:** 10.1101/834283

**Authors:** Aude Guénolé, Fabien Velilla, Aymeric Chartier, Claude Sardet, Martine Simonelig, Bijan Sobhian

## Abstract

Post-transcriptional regulatory mechanisms play a role in many biological contexts through the control of mRNA degradation, translation and localization. Here, we show that the uncharacterized RING finger protein RNF219 co-purifies and strongly associates with the CCR4-NOT complex, the major mRNA deadenylase in eukaryotes, that mediates translational repression in a deadenylase activity-dependent and -independent manner. Strikingly, although RNF219, inhibits the deadenylase activity of CCR4-NOT, it enhances its capacity to repress translation of a targeted mRNA, an effect of RNF219 that requires its interaction with CCR4-NOT. We propose that RNF219 is an interacting partner of the CCR4-NOT complex that switches the translational repressive activity of CCR4-NOT from a deadenylation-dependent to a deadenylation-independent mechanism.

## INTRODUCTION

The regulation of eukaryotic gene expression has been widely described at the transcriptional level. However, numerous studies also report the importance and the variety of post-transcriptional regulatory events. While a number of these processes such as, splicing, mRNA polyadenylation or mRNA quality control arise in the nucleus, the cytoplasmic fate of mRNAs relies mostly on mRNA stability and translation. In eukaryotes, mRNA stability and translation are intimately linked to the 5’ end cap structure and the 3’ end polyadenosine (poly(A)) tail of the mRNA. Eukaryotic mRNA decay is generally initiated by shortening of the poly(A) tail. Then, removal of the 5’ cap structure (decapping) is followed by 5’ to 3’ exonucleolytic degradation or alternatively mRNA is digested from the 3’ end (reviewed in ^1^; ^2^). Besides its function in mRNA decay, deadenylation also contributes to translation silencing ^3^. In some cases, deadenylated transcripts can be stable but untranslatable until cytoplasmic polyadenylation reactivates translation ^4^. Moreover, long poly(A) tails induce translation in oocytes and early embryos and oocytes ^3,5,6^ while in somatic cells this correlation is lost ^7^. Interestingly, a recent report from Lima et al. linked short poly(A) tails to abundant and highly translated mRNAs ^8^.

One of the major actors involved in deadenylation is the evolutionary conserved CCR4-NOT complex. The structure-function of most CCR4-NOT subunits has been well characterized in yeast, *Drosophila* and humans ^9–14^. These reports showed that CNOT1 is the largest CCR4-NOT core subunit that is required to maintain the integrity of the complex. It docks the CNOT2 and CNOT3 subunits at its C-terminus, the deadenylase subunits, CNOT7 and CNOT8, at its central MIF4G domain, and CNOT11/CNOT10 at its N-terminus ^10,11,13^.The deadenylase activity of the complex is carried out by the DEDD domain containing exonucleases CNOT7 (CAF1A) and CNOT8 (CAF1B), and the endonuclease-exonuclease-phosphatase (EEP) CNOT6 (CCR4a) and CNOT6L (CCR4b). Only one CAF1 and one CCR4 are found per CCR4-NOT complex ^14^.

CCR4-NOT plays a role in global mRNA degradation ^2,15^. Yet, it has been more specifically described for targeted mRNA decay such as nonsense-mediated mRNA decay (NMD) where CCR4-NOT is recruited by RNA binding proteins (RBPs) or miRNAs to specific 3’ untranslated regions (UTRs) in mRNAs under certain physiological conditions ^16–21^, or during developmental processes ^22,23^

Besides its role in deadenylation-dependent mRNA decay, CCR4-NOT can regulate translation in a deadenylation-indeendent manner. In this pathway, the translation repressors DDX6 (*p54/RCK)* (yeast ortholog: Dhh1) and EIF4ENIF1 (4E-T) bind to the CCR4-NOT scaffold subunit CNOT1^24,25^. These proteins can activate the decapping machinery, which leads to translation inhibition, mRNA storage and/or mRNA degradation ^26–29^.

Here, we identify the heretofore uncharacterized C3HC4 RING domain dependent E3 Ubiquitin ligase RNF219, which stably binds the CCR4-NOT complex. We further show that RNF219 inhibits the expression and stabilizes the poly(A) tail of a targeted mRNA. Moreover, loss of RNF219 interaction with CCR4-NOT greatly abolishes these two activities. Hence, we propose that RNF219 may act as a molecular switch that enables CCR4-NOT to change function from deadenylation to deadenylation-independent translational repression.

## MATERIAL AND METHODS

### Cell culture

HEK293T, HeLa, and U2OS cells were grown in DMEM (Life Technologies) containing 10% FBS (Sigma-Aldrich) and 1% penicillin/streptomycin (Life Technologies). HEK293T and HeLa RNF219 CRISPR KO (HEK293T SG1-C and HeLa SG1-C) construct and cell lines were generated using Ran et al. published protocol ^30^. The sequences used to generate the guide RNA are sgRNA_1F (CACCGCTATGCTAAGCCATACGGTC), sgRNA_1R (AAACGACCGTATGGCTTAGCATAG).

### Antibodies and reagents

Antibodies were obtained from Santa Cruz Biotechnology (Tubulin sc-5286, RPL3 sc-86828, p27 Antibody (F-8) sc-1641), Bethyl Laboratories (CNOT3 A302-156A, CNOT2 A302-562A, RNF219 A302-540A (RNF219-C)), Proteintech (CNOT1 14276-1-AP), Covance (anti-HA.11, MMS-101P) and Abcam (GAPDH ab9485). FLAG-M2 agarose (F2426 or A2220-5ML) and FLAG peptide (F3290-4MG) were purchased from Sigma-Aldrich. RNF219 specific antibodies were produced using an internal (RNF219-A) or C-terminal (RNF219-B) peptide by Abnova (Taiwan). Secondary antibodies were purchased from Cell Signaling Technology (goat anti-mouse IgG HRP-linked, #7076 and goat anti-rabbit IgG HRP-linked, #7074).

### Protein sequence alignment

Amino acid sequences were aligned with Clustal omega tool.

### E3 ubiquitin ligase activity

20 ng FH-RNF219 or FH-RNF219-CG, FLAG purified from transfected HEK293T cells under high salt conditions (0.5 M NaCl) was used in a 20 µl reaction following manufacturer instructions (BML-UW9920-0001 ENZO Life Sciences) with UBCH5a as E2 enzyme.

### Immunofluorescence

Cells grown on coverslips were washed with PBS, fixed with 3% PFA/2% sucrose in PBS for 5 minutes at room temperature (RT), washed again in PBS and subsequently incubated in cold permeabilization buffer (20 mM Tris (pH 7.4), 50 mM NaCl, 0.5% Triton X-100, 3 mM MgCl2, 0.3 M sucrose) for 5 min at RT. After washes in PBST (PBS, 0.1% Tween-20), cells were incubated 30 min at 37°C with the primary antibody diluted in PBST. Incubation with the secondary antibody was performed for 30 min at 37°C in PBST.

### Purification of RNF219-Associated Complexes and Mass Spectrometry

RNF219 complexes were purified from Dignam S100 extracts derived from 2×10^9^ HeLa S3 cells stably expressing RNF219 fused to a FLAG and an HA tag at the C-terminus (FHA-RNF219) by two-step affinity chromatography, according to a standard method ^31^. 5% of FLAG and HA immunoaffinity purified FHA-RNF219 or mock immunoprecipitations (IP) from four liters of culture were resolved on SDS-PAGE and stained with the Silverquest kit (Invitrogen). The remainder of the eluate was stained with colloidal blue (Invitrogen). Individual coomassie stained bands, or for closely migrating bands regions of the gel, were excised and subsequently analyzed by tandem mass spectrometry at the Harvard Medical School Taplin Biological Mass Spectrometry facility, Boston, MA.

### Whole-Cell Extract Preparation and Immunoprecipitation

All steps were performed at 4**°**C. β-mercaptoethanol and phenylmethylsulfonyl fluoride (PMSF) were added to cold solutions prior to use. Following harvest, cells were washed with cold PBS and then resuspended in 10 cell pellet volumes of TETN-150 buffer (20 mM Tris-HCl (pH 7.4), 0.5 mM EDTA, 0.5% Triton X-100, 150 mM NaCl, 2 mM MgCl2, 5 mM β-mercaptoethanol, 0.5 mM PMSF) and incubated for 30 min with rotation. Extracts were cleared by centrifugation at 18000 g for 20 min and transferred to fresh tubes. Antibodies were added for 3 h before 45 min incubation with protein A or G beads (Dynabeads 10002D, 10003D from Thermofisher scientific, or agarose beads sc-2001, sc-2001 from Santa Cruz Biotechnology) for rabbit or mouse IgG respectively. After washes in lysis buffer, FLAG-IP were eluted with FLAG peptide in lysis buffer. Immuno-Precipitation with other antibodies were eluted by incubation in SDS loading buffer for 3 minutes at 95°C.

### Luciferase assay

For each well of a six-well plate, HeLa and HEK293T cells were seeded 24 h prior to transfection. Cells were co-transfected with 100 ng of Renilla-5BoxB reporter ^32,33^ and 25 ng of Firefly control reporter, along with either N-terminally tagged λN-peptide and HA tagged protein expression plasmid (NHA-LacZ (25 ng), NHA-RNF219 (600 ng), NHA-RNF219-Id (900 ng), NHA-RNF219-Cd (200 ng) NHA-CNOT7 (200 ng), NHA-CNOT1-R (200 ng), FHA-RNF219 (500 ng) plasmids, using Fugene (Promega) for HeLa or calcium phosphate (Sigma) for HEK293T. Cells were harvested 24 h after transfection. Firefly and Renilla luciferase activities were analyzed individually with a specific Luciferase assay system. The Promega E151A substrate for Firefly and the Renilla-Glo Luciferase Assay System with E2720 substrate for Renilla.

### Statistical analysis

Unpaired two-tailed Student’s t-test were used to measure the statistical significance of the mean (n = 3) between conditions. Symbol meanings are: ns P > 0.05, *P ≤ 0.05, **P ≤ 0.01, ***P ≤ 0.001, ****P ≤ 0.0001.

### Proteasome inhibition assay

Renilla-BoxB (RL), FireFly (FL) plasmids and indicated N-peptide constructs were co-transfected in HEK293T cells. 16 h after transfection half of the cells were subjected to proteasome inhibitor (10 µM of MG132) treatment for 6 h. Afterwards RL activity was measured and normalized using Firefly luciferase activity.

### Quantitative RT–PCR (qRT–PCR) and poly(A) tail length analysis

Total RNA was extracted using TRIzol LS (Life Technologies). For reverse transcription, 0.5 µg of RNA was heated for 5 min to 65°C and cDNA was generated using SuperScript III (Life Technologies) and random primers for 1 h at 50°C followed by heat inactivation for 15 min at 70°C. qPCRs (for primers, see Supplemental Table S1) were performed with SYBR Green master mix (Ozyme) using a LightCycler 480 (Roche) and incubated for 2 min at 95°C with 35 cycles of 10 sec at 95°C, 25 sec at 60°C, and 25 sec at 72°C. Data were normalized to GAPDH and analyzed using the 2−ΔΔCT method ^34^. Poly(A) tail length was analyzed using the ePAT method ^35^. Briefly, 1 µg of total RNA was incubated with 5 μM oligo-(dT)-anchor (5’GCGAGCTCCGCGGCCGCGTTTTTTTTTTTT3’) and 5 U of Klenow polymerase (New England Biolabs) for 1 h at 37°C for template extension of the poly(A) tail, followed by reverse transcription using 200U of SuperScript III (Life Technologies) for 1 h at 55°C. Poly(A) tails were amplified by PCR using a gene-specific forward primer and the oligo-(dT)-anchor and migration of PCR product performed on 3% UltraPure Agarose 1000 gel (Life Technologies).

### Polyribosome purification

All steps were performed on ice or in a cold room. Dithiothreitol (DTT), RNasin and phenylmethylsulfonyl fluoride (PMSF) were added to cold solutions prior to use. Cells were treated with 1 mg/ml cycloheximide (CHX) 5 min prior harvesting. Following harvest, cells were washed with cold PBS containing 100 µg/ml CHX and resuspended in lysis polysome buffer A (20 mM Tris-HCl (pH 7.5), 100 mM KCl, 10 mM MgCl2, 0.2 mg/ml heparin, 100 µg/ml CHX, 1% Triton, 1 mM DTT, 2.5 mM PMSF, 6U/ml RNasin) and incubated 20 min with rotation.

After 6 min centrifugation at 18000 g, 1 ml of extract was loaded onto a 11 ml 10% to 50% sucrose gradient in buffer B (20 mM Tris-HCl (pH 7.5), 50 mM KCl, 10 mM MgCl2, 100 µg/ml CHX, 1 mM DTT, 2.5 mM PMSF, 6U/ml RNasin) and centrifuged for 2 h at 33000 rpm in a Beckman SW41 rotor. 750 µL fractions were collected from the top of the gradient. RNA was extracted with Trizol LS reagent after a proteinase K treatment. RNA samples were treated for 1h at 37 °C with DNase I (1U per mg of RNA; Ambion), extracted again with Trizol LS and precipitated.

Alternatively, 200 µl extract was loaded onto a 4.5ml sucrose gradient and spun in a SW55Ti rotor at 46 000 rpm for 70 min. Fractions were manually collected from the top of the gradient. 1% SDS was added to fractions of interest prior to RNA extraction with acidic phenol chloroform (Ambion 9720) following manufacturer instructions.

### RNA-seq procedure and sequences analysis

HeLa cells were transfected with either a control siRNA (siSCR) or a siRNA against RNF219 (CDS or UTR2). RNA was extracted with Trizol LS reagent according to the manufacturer’s instructions. RNA samples were then treated with DNase I, extracted again with Trizol LS and precipitated.

Ribosomal RNA depletion and RNA-Seq library preparation were performed at Fasteris SA (Plan-les-Ouates, Switzerland) using TruSeq stranded mRNA kit (Illumina). RNAseq samples were sequenced using Illumina HiSeq High-Output (HO), single-end, 50 bp reads at Fasteris SA.

RNA-seq files were analysed at Fasteris SA. Briefly, RNA-Seq files were aligned to the reference human genome (hg38; https://my.illumina.com/Message/iGenome/) using Bowtie 2 (http://sourceforge.net/projects/bowtie-bio/files/bowtie2/2.0.5/). Tophat was used as the splice junction mapper for RNA-Seq reads (http://tophat.cbcb.umd.edu/). Samtools was used as the toolbox for manipulation of SAM/BAM files.

Cufflinks was used to assemble aligned RNA-Seq reads into transcripts, estimate their abundances, and test for differential expression and regulation transcriptome-wide (http://cufflinks.cbcb.umd.edu/). CummeRbund was used for the exploration, analysis and visualization of Cufflinks output (http://compbio.mit.edu/cummeRbund/).

GO term analysis was done using PANTHER Overrepresentation Test (which can be found on the geneontology.org website).

### Data availability

RNA-sequencing data have been deposited in GEO under the accession number GSE95442.

## RESULTS

### Characterization of RNF219: an unstudied RING dependent ubiquitin ligase

We initially identified RNF219 as a substoichiometric interactor of the HIV-1 Tat transcription factor. RNF219 was found in the list of proteins co-purifying with Tat that were identified by mass-spectrometry in the study by Sobhian et al. ^36^ (Personal communication). RNF219 is a 726 amino acid protein of unknown function containing at its N-terminus a 37-amino acid length C3HC4 RING finger domain (Fig. 1A, 1B) ^37,38^. RNF219 orthologs can be identified among most vertebrates, but not among other metazoans (Fig. 1B). Immuno-localization of FLAG-HA tagged RNF219 (FHA-RNF219) expressed in U2OS cells shows that this protein is found in both the cytoplasm (60% of cells n=110, Fig. S1A) and the nucleus (40% of cells n=110, Fig. S1A). Of note, RNF219 was not found in both compartments simultaneously in the scored nuclei.

**Figure 1.**
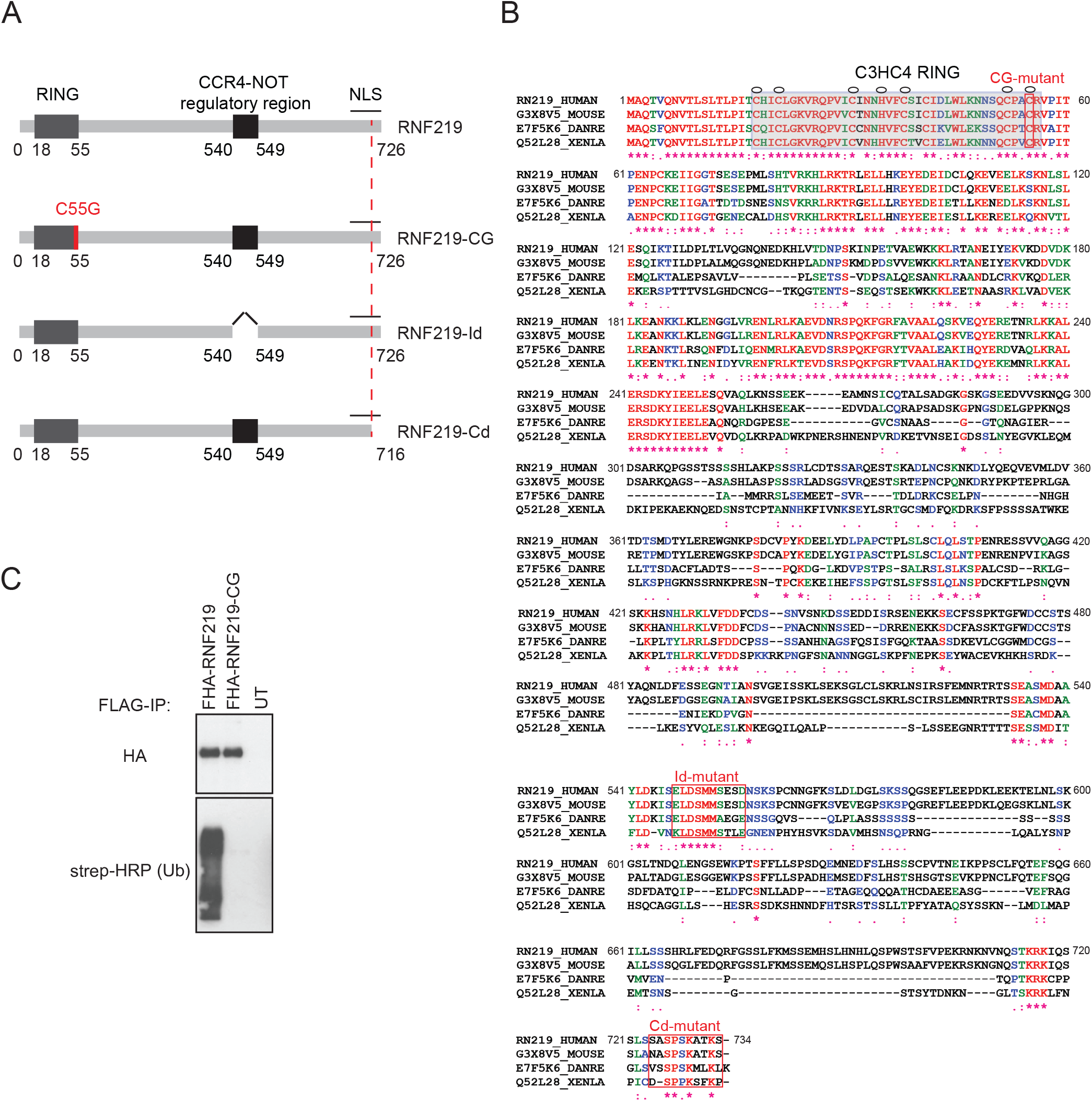
RNF219 is a RING dependent ubiquitin ligase. (A) Schematic representation of the RNF219 protein and mutants used in this study. (B) Sequence alignment of the human hRNF219 protein and its *Mus musculus*, *Xenopus laevis* and *Danio rerio*, orthologs. Amino acid sequences were aligned with the Prabi CLUSTALW tool. Identical residues are annotated with an asterisk (*) and coloured in red, whereas similar residues are annotated with two (:) and coloured in green and less similar residues are annotated with one dot (.) and coloured in blue. One conserved RING domain of C3HC4 type was predicted in the N-terminus of all the conserved alignments (Highlighted in grey); Zinc coordinating residues are marked with a circle. Mutated residues and corresponding mutants are highlighted by a red box. (C) RNF219 but not the RING mutant can form poly-ubiquitin chains in an in vitro assay. FHA-RNF219 and FHA-RNF219-CG (a RNF219 RING mutant where the cysteine 55 was replaced by a glycine) affinity purified with FLAG antibody and peptide eluted from transfected 293T cells, were incubated with E1 enzyme, UBCH5a, and biotinylated ubiquitin (lanes 1 and 2, respectively). The third lane contains the purification from untransfected cells as negative control. Ubiquitin was detected with HRP conjugated streptavidin (Strep-HRP).

To better characterize RNF219, we generated two polyclonal antibodies (αRNF219-A and αRNF219-B) directed against human RNF219, and two RNF219 CRISPR KO cell lines (HEK293T SG1-C and HeLa SG1-C) (Fig. S1B-C). Immunoblots performed on HeLa cell extracts showed that αRNF219-A specifically recognizes a 85 kDa protein that disappeared upon RNF219 depletion using two different small interfering RNAs (siRNAs), targeting RNF219 coding sequence (CDS) or 3’UTR (UTR), respectively (Fig. S1B). The second antibody (αRNF219-B) was used to immuno-precipitate endogenous RNF219 (Fig. S1D, 2C, S2A, S2B). αRNF219-C antibody (from Bethyl laboratory) was used to immuno-precipitate endogenous RNF219 in order to confirm the data obtained with αRNF219-B (Fig. S2B).

Since RNF219 contains a RING finger domain (Fig. 1B), found in many E3 ubiquitin ligases, we tested its ability to catalyse the formation of polyubiquitin chains. An in vitro E3 ubiquitin ligase assay was performed with FHA-RNF219, immunoprecipitated from 293T cell extract under high salt conditions, in the presence of E1, E2 (UBCH5a) enzymes and biotinylated ubiquitin. Wild-type FHA-RNF219, but not a mutant form containing a point mutation in the RING domain (cysteine 55 changed to a glycine; RNF219-CG), was able to form polyubiquitin chains in this biochemical assay (Fig. 1C). This result suggests that RNF219 is a RING dependent ubiquitin ligase.

### RNF219 associates with the CCR4-NOT complex

We purified RNF219 protein complexes to address its cellular function. TAP-tag experiments ^31^ were performed using FLAG/HA-tagged RNF219 (FHA-RNF219) stably expressed in HeLa S3 cells. Immunoprecipitated material was analysed by tandem mass spectrometry. Remarkably, all subunits of the CCR4-NOT except CNOT4 were identified with the highest peptide coverage (Fig. 2A and highlighted in yellow in Table S2). Of note, we did not observe major differences in the peptide coverage measured for the different CCR4-NOT deadenylase subunits CNOT6, CNOT6L, CNOT7 and CNOT8, suggesting that RNF219 can stably associate with CCR4-NOT complexes containing either subunit.

**Figure 2.**
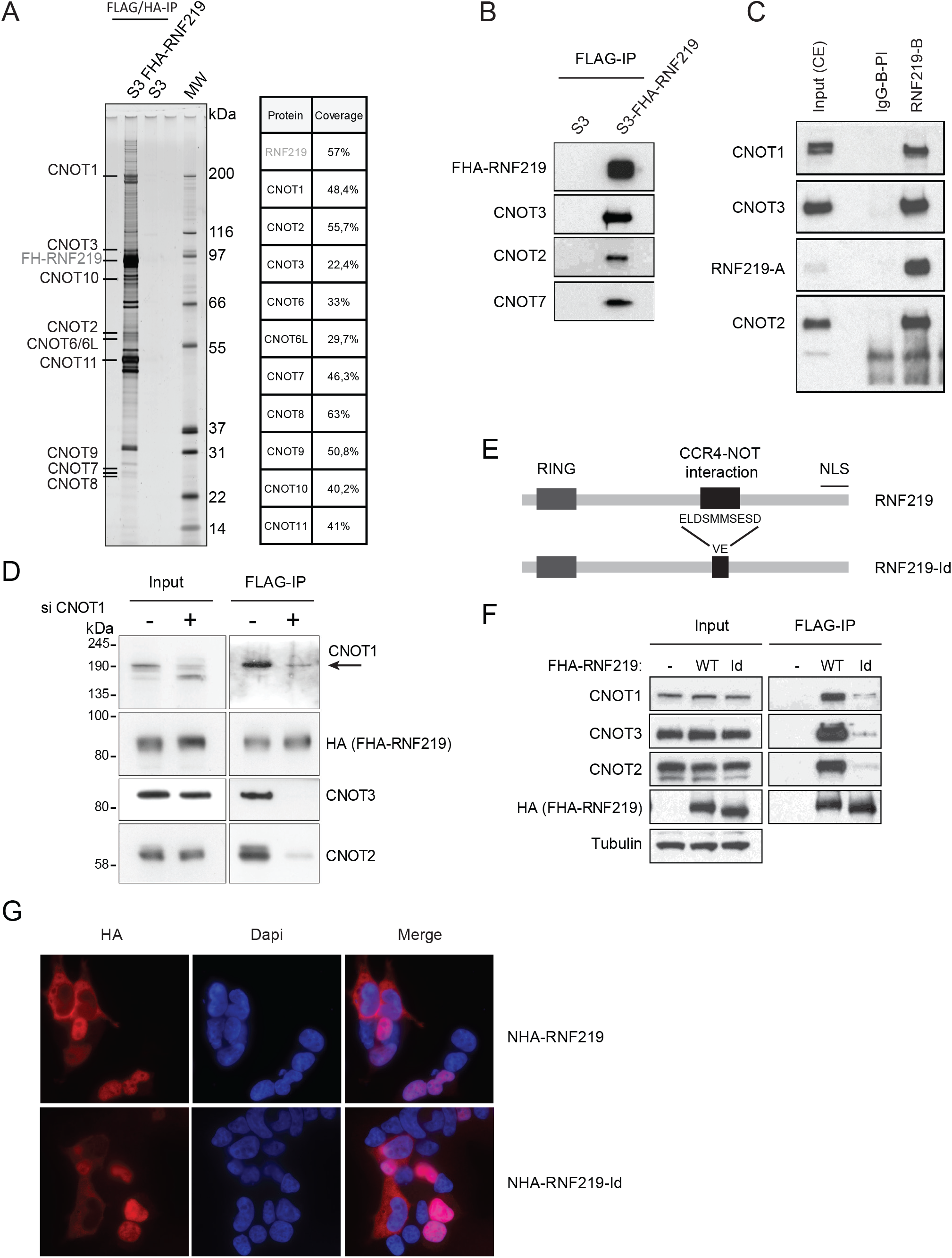
RNF219 binds to the CCR4-NOT complex. (A) (Left) FHA-RNF219 was purified from Dignam cytoplasmic S100 extracts prepared from stably transduced HeLa S3 cells (S3-FHA-RNF219) or non-transduced HeLa S3. FHA-RNF219 was sequentially purified on anti-FLAG and anti-HA antibody-conjugated agarose beads. Proteins were resolved by SDS-PAGE and visualized by silver staining. (Right) The identity of FHA-RNF219-associated proteins was determined by tandem mass spectrometry and percent peptide coverage by amino acid count for CCR4-NOT subunits detected are listed. (B) FLAG-immuno-precipitations from S3-FHA-RNF219 cell extracts confirm that FHA-RNF219 is associated with a number of CCR4-NOT subunits as determined by mass spectrometry. The presence of CCR4-NOT subunits CNOT3, CNOT2, CNOT7 in the IPs was analyzed by immunoblotting (IB). (C) Endogenous RNF219 was immuno-purified from HeLa cell extracts using a specific antibody against RNF219 (RNF219-B) or pre-immune IgG control (IgG-B-PI). The presence of the CCR4-NOT subunit CNOT1, CNOT2 and CNOT3 in the IP was analyzed by IB. RNF219 was detected with our RNF219-A antibody (long exposure is shown in Fig. S2B). (D) RNF219/CNOT3, RNF219/CNOT2 interactions are CNOT1 dependent. FHA-RNF219 expressing HeLa cells were transfected with siRNA against CNOT1 (+) or control siRNA (-). IPs were performed using anti-FLAG agarose beads and were blotted for the indicated proteins. The arrow indicates the expected CNOT1 band. The lower is a non-specific band as it does not disappear in siCNOT1 treated cells. (E) Schematic representation of RNF219-Id, a CCR4-NOT complex interaction defective allele. Amino acids 540 to 549 (ELDSMMSESD) were replaced by VE. (F) RNF219-Id interacts with the CCR4-NOT complex very inefficiently. IPs were performed using anti-FLAG agarose beads and were immunoblotted for the indicated proteins. (G) RNF219-Id localizes both in the nucleus and in the cytoplasm similarly to WT RNF219. Immuno-fluorescence was performed with anti-HA antibody on HEK293T cells transfected with the indicated constructs.

CNOT3, CNOT2 and CNOT7 co-purification with FHA-RNF219 were confirmed by regular RNF219-FLAG immunoprecipitation followed by immunoblotting with antibodies directed against these CCR4-NOT subunits (Fig. 2B). Importantly, endogenous RNF219 co-immunoprecipitated endogenous CCR4-NOT subunits using two different antibodies (RNF219-B and RNF219-C) for IP (Fig. 2C, S2A and S2B). To control that the interaction is not due to a protein non-specifically pulled down with RNF219-B antibody, immunoprecipitation was performed with RNF219-B antibody on RNF219-Crispr knockout cells transfected or not with a plasmid containing RNF219 cDNA (Fig. S1D; respectively WT and Mock). This experiment shows that CNOT2 co-immunoprecipitates with RNF219-B antibody only in RNF219 knockout cells reconstituted with WT RNF219 (Fig S1D) which confirms the specificity of RNF219 interaction with at least one subunit of the complex.

As CNOT1 is the core subunit of the CCR4-NOT complex ^9–11,39^, we next hypothesized that the depletion of CNOT1 could abrogate RNF219 association with subunits of the complex. As predicted, upon siRNA-mediated depletion of the CNOT1 scaffold, RNF219 co-precipitation with the other components of the CCR4-NOT complex, CNOT2 and CNOT3, is very much reduced (Fig. 2D).

To identify the CCR4-NOT interaction domain, we truncated RNF219 protein in multiple fragments until we observed that CCR4-NOT interaction was lost (Fig. S2C-D). RNF219 was first truncated in six large fragments with a truncation step of 121 amino acids (represented by F1 to F6 in Fig S2C). The F5 fragment failed to interact with at least three CCR4-NOT subunits (CNOT1, CNOT2, CNOT3; Fig. S2C). That F5 fragment was then truncated in four fragments with a truncation step of 10 amino acids (represented by F5F, F5G, F5H, F5I in Fig. S2D). CCR4-NOT interaction was lost with the F5G fragment (Fig. S2D). This led to the identification of a minimal domain necessary for RNF219 binding to the CCR4-NOT complex (Fig. 2E-F; Fig. S2D). Replacement of the amino acids ELDSMMSESD by VE in the middle part of RNF219 strongly reduces RNF219 interaction with CNOT1, CNOT2 and CNOT3 (Fig. 2F). We named this interaction mutant RNF219-Id. Of note, the interaction between RNF219 and the CCR4-NOT complex is RNAse resistant under conditions dissociating the 7SKsnRNP (^40,41^ and Fig. S2E) indicating that the CCR4-NOT and RNF219 interaction is not mediated by RNA. However, RNA may play a role during the formation of the complex. Finally, we showed by immuno-localization that RNF219-Id, similarly to WT RNF219, is present either in the nucleus or in the cytoplasm (Fig. 2G).

Altogether, our data identify RNF219 as a new CCR4-NOT interacting protein, requiring the CNOT1 scaffold but not RNA for its association to the complex.

### RNF219 represses the expression of a targeted mRNA

To characterize RNF219 functionally, we asked whether RNF219 plays a role in post-transcriptional regulation of mRNA, a well-established CCR4-NOT function. We set up a classical λNpeptide/BoxB RNA tethering reporter assay to address this question (described and characterized in: ^32,33,42,43^). Briefly, in this assay, a λNpeptide-tagged candidate protein is targeted to a Renilla luciferase (RL) reporter mRNA through binding of the λNpeptide to BoxB sites present in the reporter 3’UTR (Fig. 3A).

**Figure 3.**
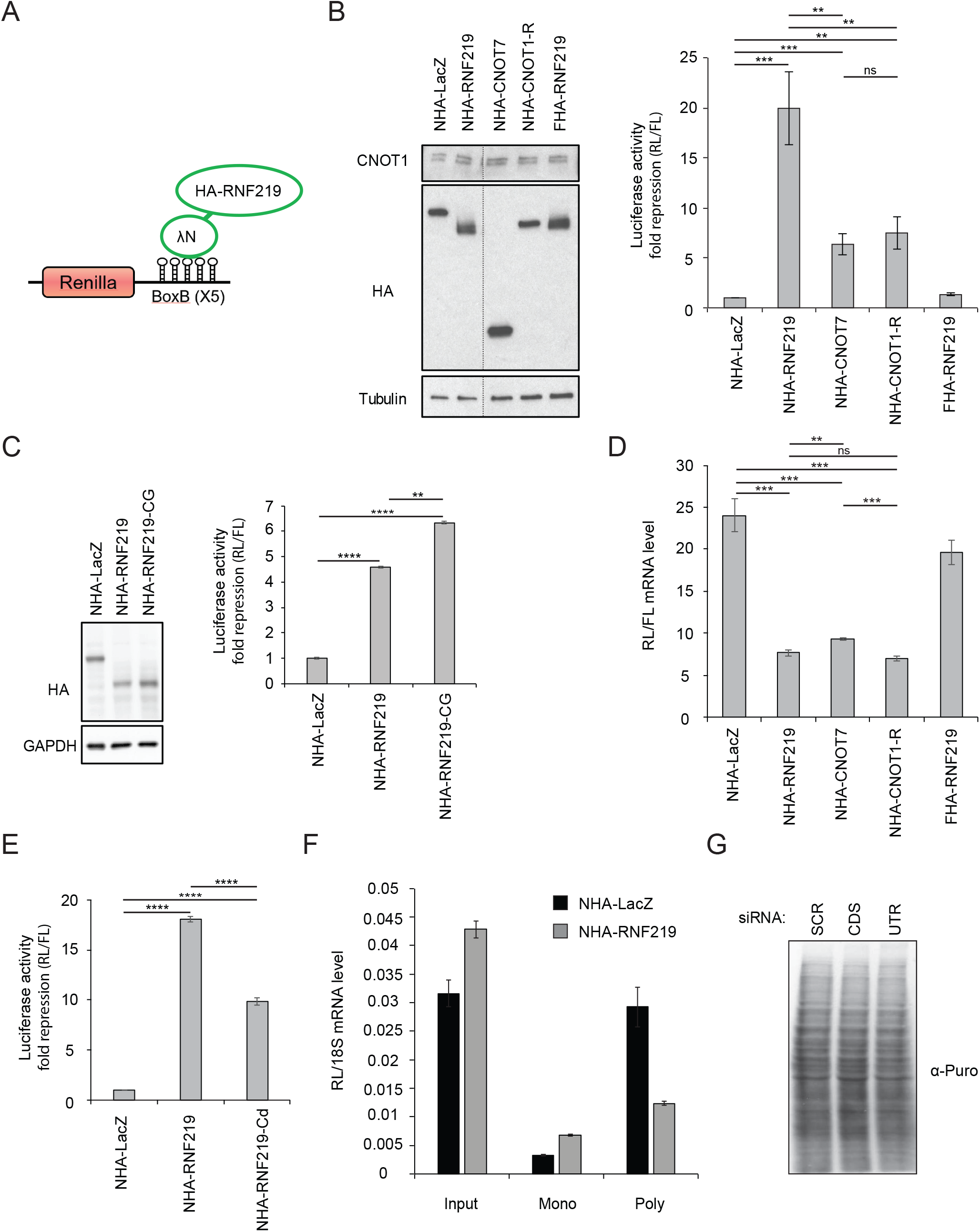
RNF219 affects the translation of a targeted mRNA. (A) Schematic representation of the Renilla Luciferase (RL) reporter mRNA, containing the Renilla luciferase gene and five 19-nt BoxB hairpins sequences at its 3’UTR. Recruitment of HA-tagged protein to the 3’UTR is mediated by the fused λN-peptide, which has a high affinity for the BoxB sequence. (B) Recruitment of RNF219 to the 3’UTR mRNA of the RL reporter inhibits its expression. RL activity was determined in the indicated conditions (NHA-LacZ, NHA-RNF219, NHA-CNOT7, NHA-CNOT1-R, FHA-RNF219). Renilla Luciferase (RL) activity was normalized on Firefly luciferase (FL) activity expressed from a plasmid not containing BoxB sequences, co-transfected with that of Renilla. RL repression levels are relative to that of the control NHA-LacZ set to 1. Error bars represent SD, n = 3. **P < 0.01, ***P < 0.001 based on unpaired two-tailed Student’s t-test. Protein levels of NHA-LacZ, NHA-RNF219, NHA-CNOT7, NHA-CNOT1-R, FHA-RNF219 in HeLa cells transfected with the corresponding plasmids were analyzed by immunoblotting. (C) Ubiquitin ligase activity of RNF219 is not necessary for its repressive role. RL activity was determined in the indicated conditions (NHA-LacZ, NHA-RNF219, NHA-RNF219-CG). Renilla Luciferase (RL) activity was normalized as in (B). Error bars represent SD, n = 3. **P < 0.01, ****P < 0.0001 based on unpaired two-tailed Student’s t-test. Protein levels of NHA-LacZ, NHA-RNF219, NHA-RNF219-CG, in HeLa cells transfected with the corresponding plasmids were analyzed by immunoblotting. (D) Recruitment of RNF219 to the 3’UTR mRNA of the RL reporter leads to decreased reporter mRNA level. Total RNA from the indicated conditions (NHA-LacZ, NHA-RNF219, NHA-CNOT7, NHA-CNOT1-R, FHA-RNF219) was extracted as described in the methods. RT-QPCR was performed using primers specific from both RL and FL reporters to quantify their respective mRNA level (the RT-QPCR primers used are indicated in table S1). RL mRNA level was normalized on FL mRNA level. Error bars represent SD, n = 3. **P < 0.01, ***P < 0.001, based on unpaired two-tailed Student’s t-test. (E) RNF219 mutant (RNF219-Cd) that resides predominantly in the cytoplasm also represses the RL expression. RL activity was determined in the indicated conditions (NHA-LacZ, NHA-RNF219, NHA-RNF219-Cd). Renilla Luciferase (RL) activity was normalized as in (B). Error bars represent SD, n = 3. ****P < 0.0001 based on unpaired two-tailed Student’s t-test. (F) RL mRNA level in the monoribosomal (Mono) and polyribosomal fractions (Poly, fractions 10-11 on Fig. S3C) of extracts from HeLa cells transfected with NHA-LacZ or NHA-RNF219. Polyribosome purification was done using the alternative protocol described in the methods. mRNA levels were quantified by RT-QPCR and were normalized to 18s mRNA levels. (G) RNF219 does not affect global translation. 24h after siRNA treatment (two different siRNA against RNF219: CDS & UTR; Fig. S1B) a 30-minute puromycin pulse is analysed by immunoblot using an anti-puromycin antibody. The two siRNA against RNF219 show no difference in global puromycin incorporation compare to the control (SCR).

HEK293T cells were co-transfected with the RL reporter and either λN and HA-tagged-LacZ (control NHA-LacZ), or -RNF219 construct (NHA-RNF219; Fig 3B, C, E Fig. S3A-B). Subsequently, the RL reporter expression was assessed by measuring the luciferase activity. Interestingly, tethering RNF219 strongly repressed the RL reporter mRNA expression (20-fold compared to control; Fig. 3B). RNF219-mediated repression was even higher compared to the two CCR4-NOT subunit CNOT7 and the CNOT1-R fragment ^24^(6- and 8-fold for NHA-CNOT7 and NHA-CNOT1-R respectively; Fig. 3B) for which repression had already been described ^24,26,44,45^. Of note, FHA-RNF219 not fused to λN-peptide did not repress the reporter (Fig. 3B, last bar), suggesting that NHA-RNF219 acts in cis.

Next, we wondered whether the ubiquitin-ligase activity was necessary for RNF219 to repress the RL reporter. Hence, we tested the RING-mutated version of RNF219 described in Figure 1C, in the tethering assay. Surprisingly, RNF219-CG was able to repress the reporter similarly to WT RNF219 (Fig 3C).

This suggests either that the ubiquitin ligase activity of RNF219 is not necessary for its repressive role or, more likely, that the ubiquitin ligase activity functions at steps bypassed in the assay.

### RNF219-mediated repression is not solely due to mRNA or protein degradation

CCR4-NOT tethering to reporter mRNA classically leads to RNA degradation ^46,47^. Thus, we monitored the RL reporter mRNA level in these cells to test whether repression of luciferase activity was due to a decrease in mRNA quantity (Fig. 3D). Expectedly, tethering CNOT7 or CNOT1-R induced a 2 to 2.5-fold decrease in mRNA level. Tethering RNF219 also led to a 2.5-fold decrease in mRNA level, however the repression of the expression of the reporter was much stronger with RNF219 than with CNOT7 or CNOT1-R. This suggests that mRNA degradation alone does not explain the full RNF219-mediated RL reporter repression.

Next, we tested whether the decreased reporter expression level was a consequence of protein degradation. However, inhibiting protein degradation using a proteasome inhibitor (MG132) did not rescue the RNF219 mediated RL repression (Fig. S4A). This indicates that RNF219 does not affect RL reporter expression through protein degradation.

We observed that RNF219 localizes either in the nucleus or in the cytoplasm (Fig. S1A). Thus to question the importance of RNF219 localisation for its repressive activity, we tested a RNF219 mutant (RNF219-Cd) that resides predominantly in the cytoplasm (Fig. S4B). Importantly, we found that RNF219-Cd also represses the RL expression indicating that this activity is primarily a cytoplasmic process (Fig. 3E) and could not be explained by the sequestration of mRNA in the nucleus as only mechanism.

### RNF219 inhibits mRNA translation

To further explore the molecular mechanism by which RNF219 represses the RL mRNA expression, we asked whether RNF219 affects mRNA translation. To answer this question, we purified poly-ribosomes from cells transfected with the RL reporter and NHA-LacZ or NHA-RNF219 constructs. Mono and poly-ribosomal fractions were identified by high level of RPL3, a ribosomal subunit (Fig. S4C). We arbitrarily took two poly-ribosomal fractions out of the three we purified (fractions 10 and 11 in Fig. S4C) and quantify the mRNA level of the RL reporter. 18s mRNA level was used as internal control. RL reporter mRNA was significantly less abundant in the poly-ribosomal fraction of cells expressing NHA-RNF219 compared to cells expressing NHA-LacZ (Fig. 3F; Fig. S4D). To control that RNF219 did not affect global translation, puromycin incorporation in control (siSCR) or RNF219 (siCDS and siUTR) depleted cells was measured (^48^; Fig. 3G; Fig. S4E, S1B). No difference was detected in global translation activity between both conditions, suggesting that RNF219 is involved in translational repression of specific mRNAs.

### RNF219-mediated mRNA translation repression is CCR4-NOT-dependent

To determine if RNF219 inhibition capacity was CCR4-NOT-dependent, we first verified that NHA-RNF219 recruited CCR4-NOT to the reporter mRNA in the tethering assay (Fig. 4A). Endogenous CNOT3 immunoprecipitation showed that CNOT3 specifically co-purified with the reporter mRNA, in presence of NHA-RNF219 compared to NHA-LacZ, consistent with the recruitment of CCR4-NOT to the mRNA by NHA-RNF219 (Fig. 4A).

**Figure 4.**
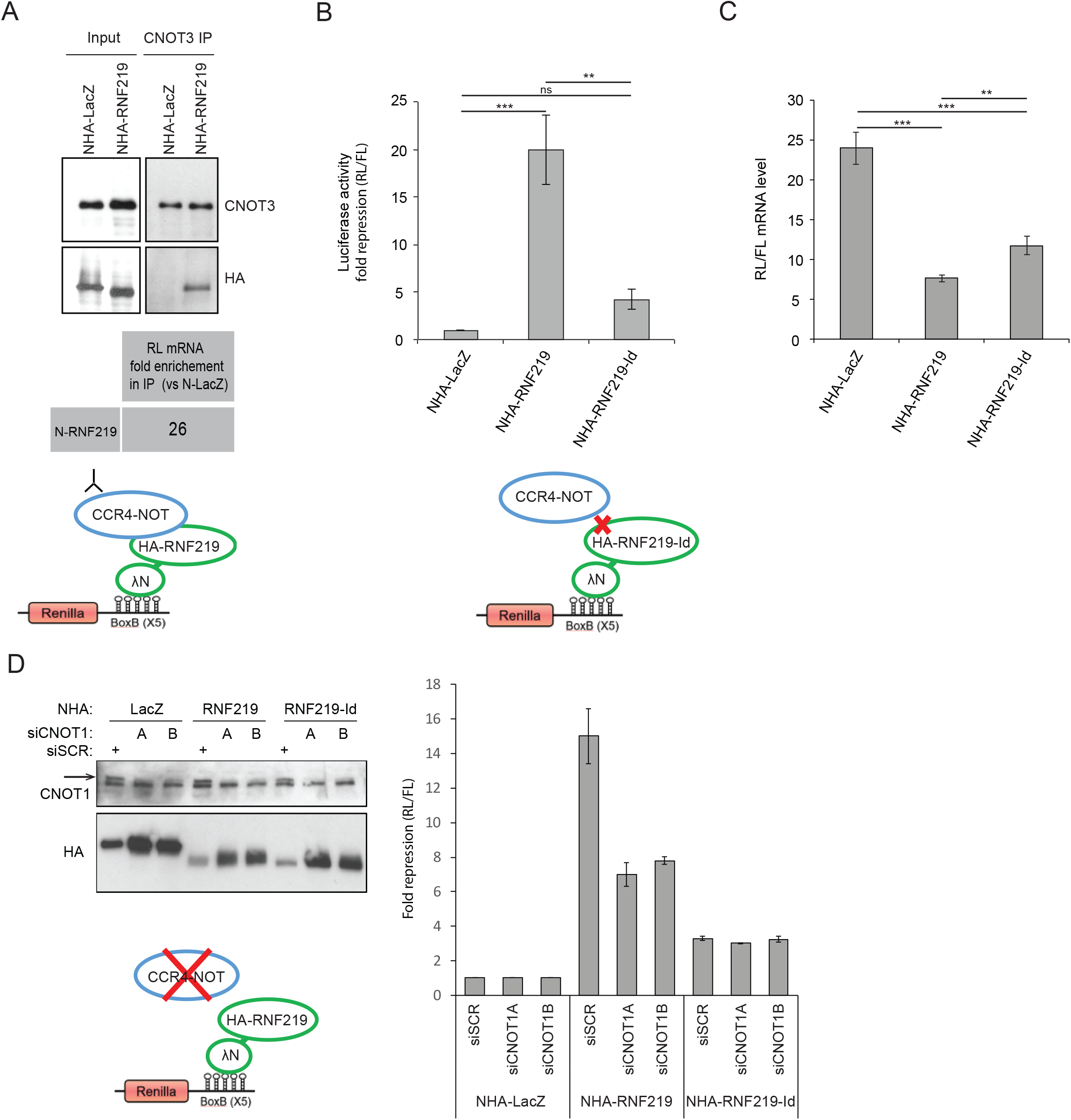
RNF219 mediated translational repression is CCR4-NOT dependent. (A) CNOT3 interacts with RNF219 tethered to the mRNA reporter. RNA-IPs were performed on extracts of cells transfected with the RL reporter and NHA-LacZ or NHA-RNF219 using CNOT3 antibody. The level of Renilla mRNA in the CNOT3-IP was highly enriched in NHA-RNF219 transfected cells as compared to NHA-LacZ transfected cells. (B) RNF219 interaction with the CCR4-NOT complex is necessary for RFN219 to repress RL reporter translation efficiently. RL activity was determined in the indicated conditions (NHA-LacZ, NHA-RNF219 and NHA-RNF219-Id). RL activity repression was determined as in Figure 3B. Error bars represent SD, n = 3. **P < 0.01, ***P < 0.001 based on unpaired two-tailed Student’s t-test. (C) mRNA levels of samples in (B) were quantified as in Figure 3D. Error bars represent SD, n = 3. *P < 0.05, **P < 0.01, based on unpaired two-tailed Student’s t-test. (D) Knock down of CCR4-NOT scaffold CNOT1 affects NHA-RNF219 mediated repression. RL activity was determined in the indicated conditions (NHA-LacZ, NHA-RNF219 and NHA-RNF219-Id) in control cells (siSCR) or in siCNOT1 depleted cells (siCNOT1A and siCNOT1B are two siRNA targeting different regions of CNOT1 mRNA). RL activity repression was determined as in Figure 3B. The arrow indicates the expected CNOT1 band. The lower is a non-specific band as it does not disappear in siCNOT1 treated cells.

Second, we tested the CCR4-NOT interaction defective allele (NHA-RNF219-Id) in the tethering assay. This mutant was four-fold less efficient in repressing the RL reporter (Fig. 4B; 20-fold for RNF219 compared to less than 5-fold for RNF219-Id). Of note, even though mRNA level was slightly higher in NHA-RNF219-Id than in NHA-RNF219 (Fig. 4C), it could not explain the lower repression observed with NHA-RNF219-Id (Fig. 4B). This implies that RNF219 acts at least partially through its interaction with CCR4-NOT to repress the RL reporter mRNA translation. We further confirmed this data, using siRNA against CNOT1 (siCNOT1A and siCNOT1B) that abolished RNF219 association with CCR4-NOT (Fig. 2D). Consistently, the RNF219 mediated translational repression of the reporter mRNA was reduced by 2-fold in CNOT1-depleted cells (Fig. 4D). Interestingly, CNOT1 depletion did not further reduce the residual repressive activity of RNF219-Id suggesting that RNF219 may also have a role in translational repression that is independent of CCR4-NOT.

These results indicate that RNF219 in complex with CCR4-NOT represses mRNA translation.

### RNF219 affects the poly(A) tail length of a targeted mRNA

The CCR4-NOT complex can repress translation either through its associated deadenylation activity or through the DDX6 pathway acting at the 5’ end of mRNA ^24,25^. Several CCR4-NOT associated E3 ligases have been reported to increase deadenylation activity leading to accelerated mRNA decay ^47,49^. We therefore analysed the poly(A) tail length of RNF219-tethered reporter mRNA using the extension Poly(A) Test (ePAT) assay ^35,50^. As expected, tethering the CCR4-NOT subunit CNOT7 resulted in shortening of the RL reporter mRNA poly(A) tail which can be observed by the accumulation of signal at the bottom of the lanes (Fig. 5A; lane 4). RNF219 tethering had the opposite effect and led to an increased poly(A) tail length of the reporter mRNA (Fig. 5A, compare lane 1 and 2). As a control, the poly(A) tail of endogenous GAPDH mRNA, not targeted by lambda peptide, was not affected.

**Figure 5.**
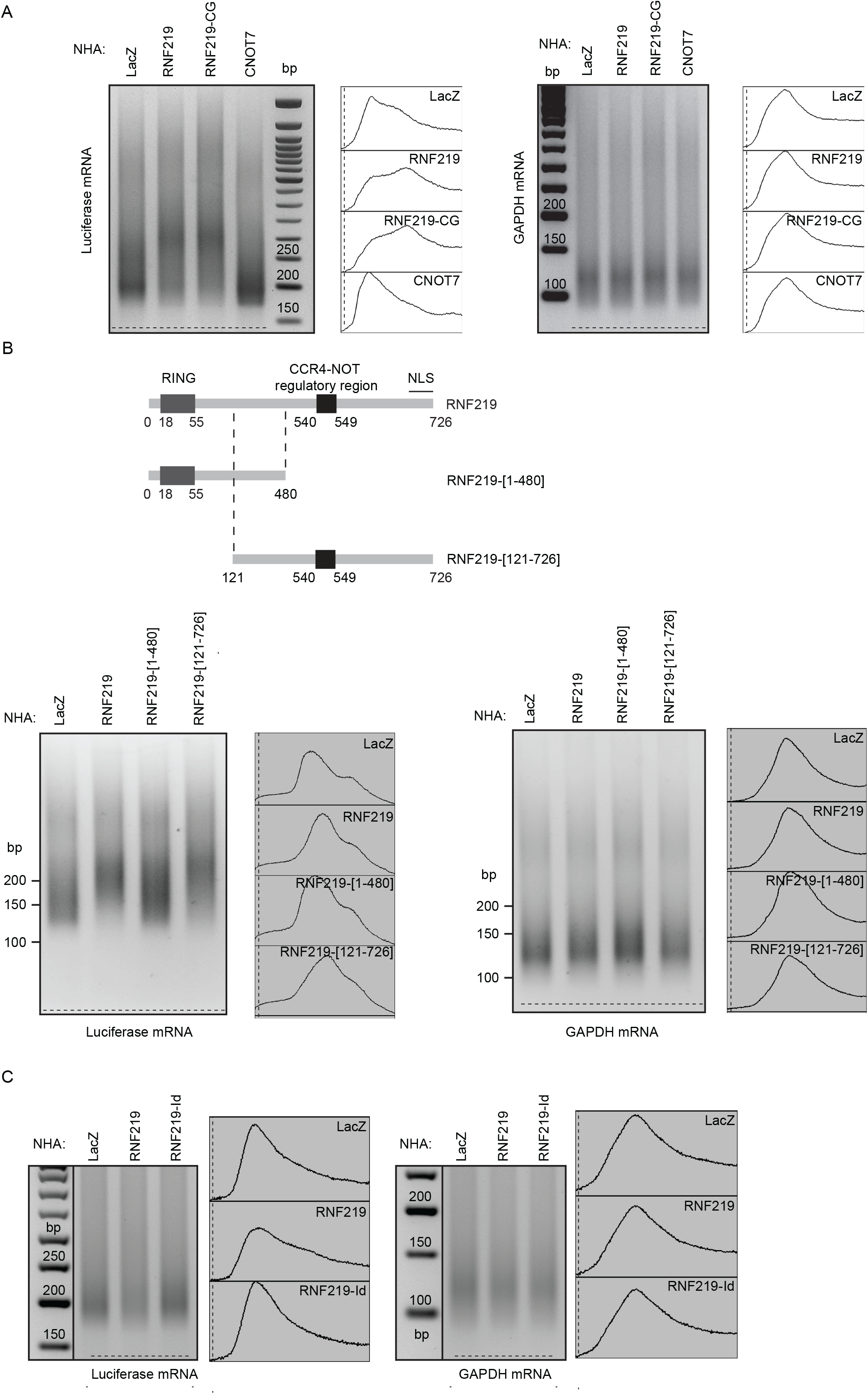
RNF219 affects the polyA tail length of a targeted mRNA. (A) RNF219 stabilizes the reporter mRNA poly(A) tail. ePAT assays were performed using HeLa cells expressing the RL reporter and either of the following constructs: NHA-LacZ, NHA-RNF219, NHA-RNF219-CG or NHA-CNOT7. On the left panel, a PAT primer (Luc_3; Table S1) specific of the poly(A) tail of the RL reporter gene is used. On the right panel a PAT primer specific of the poly(A) tail of the GAPDH control gene is used (Table S1). Corresponding image analyzer profiles are shown at the right of each ePAT gel. (B) RNF219 mRNA poly(A) tail stabilization does not depend on its RING Finger domain but depends on its interaction with the CCR4-NOT complex. ePAT assays were performed using HeLa cells expressing the RL reporter and either of the following constructs: NHA-LacZ, NHA-RNF219, NHA-RNF219-[1-480], NHA-RNF219-[121-726]. On the left panel, a PAT primer (Luc_2; Table S1) specific of the poly(A) tail of the RL reporter gene is used. On the right panel a PAT primer specific of the poly(A) tail of the GAPDH control gene is used. Corresponding image analyzer profiles are shown at the right of each ePAT gel. (C) RNF219 mRNA poly(A) tail stabilization depends on its interaction with the CCR4-NOT complex. ePAT assays were performed using HeLa cells expressing the RL reporter and either of the following constructs: NHA-LacZ, NHA-RNF219, NHA-RNF219-Id. On the left panel, a PAT primer (Luc_3; Table S1) specific of the poly(A) tail of the RL reporter gene is used. On the right panel a PAT primer specific of the poly(A) tail of the GAPDH control gene is used. Corresponding image analyzer profiles are shown at the right of each ePAT gel.

We further hypothesized that this effect was dependant on RNF219 interaction with CCR4-NOT. An RNF219 interaction mutant NHA-RNF219-[1-480] (upper panel Fig. 5B and Fragment F5 in Fig. S2C), which lacks the C-terminus containing the interaction domain and is unable to repress the RL reporter expression (Fig. S4A-B) was tested for its effect on the reporter mRNA poly(A) tail length (Fig. 5B). While RNF219 lengthened it, the interaction mutant NHA-RNF219-[1-480] did not increase the reporter mRNA poly(A) tail (Fig. 5B). We also tested the NHA-RNF219-Id interaction defective allele, unable to recruit the CCR4-NOT complex (Fig. 2E-F). This mutant did not cause lengthening of the poly(A) tail of the reporter (Fig. 5C, compare lane 3 with lane 2). These results suggest that RNF219 interaction with CCR4-NOT is necessary for RNF219-mediated inhibition of mRNA deadenylation.

Of note, NHA-RNF219-CG, the RING domain mutant (Fig. 1C) as well as a RNF219 RING-depleted allele, NHA-RNF219-[121-726], displayed similar activity as WT RNF219, indicating that the ubiquitin ligase activity is not necessary for RNF219 role in mRNA poly(A) tail lengthening in our tethering assay (Fig. 5A; lane 3, Fig. 5B; lane 4). As we previously commented (Fig. 3C), it is likely that RNF219 ubiquitin ligase activity serves at steps bypassed in the assay.

### Endogenous RNF219 has a role in cell cycle regulation

To explore the in vivo function of RNF219, we asked whether RNF219 affects the expression of p27, a known target of CCR4-NOT and member of the Cip/Kip family of cyclin dependent kinase inhibitor that triggers G1 cell cycle arrest ^51^. Depletion of RNF219 using two different siRNA (CDS and UTR), triggers an increase of p27 protein level similar to that observed in CNOT3 depleted cells (Fig. 6A). Notably, p27/CDKN1b mRNA level in RNF219 depleted cells were unchanged compared to control cells (Fig. 6B), suggesting that the effect of RNF219 on p27 is post-transcriptional.

**Figure 6.**
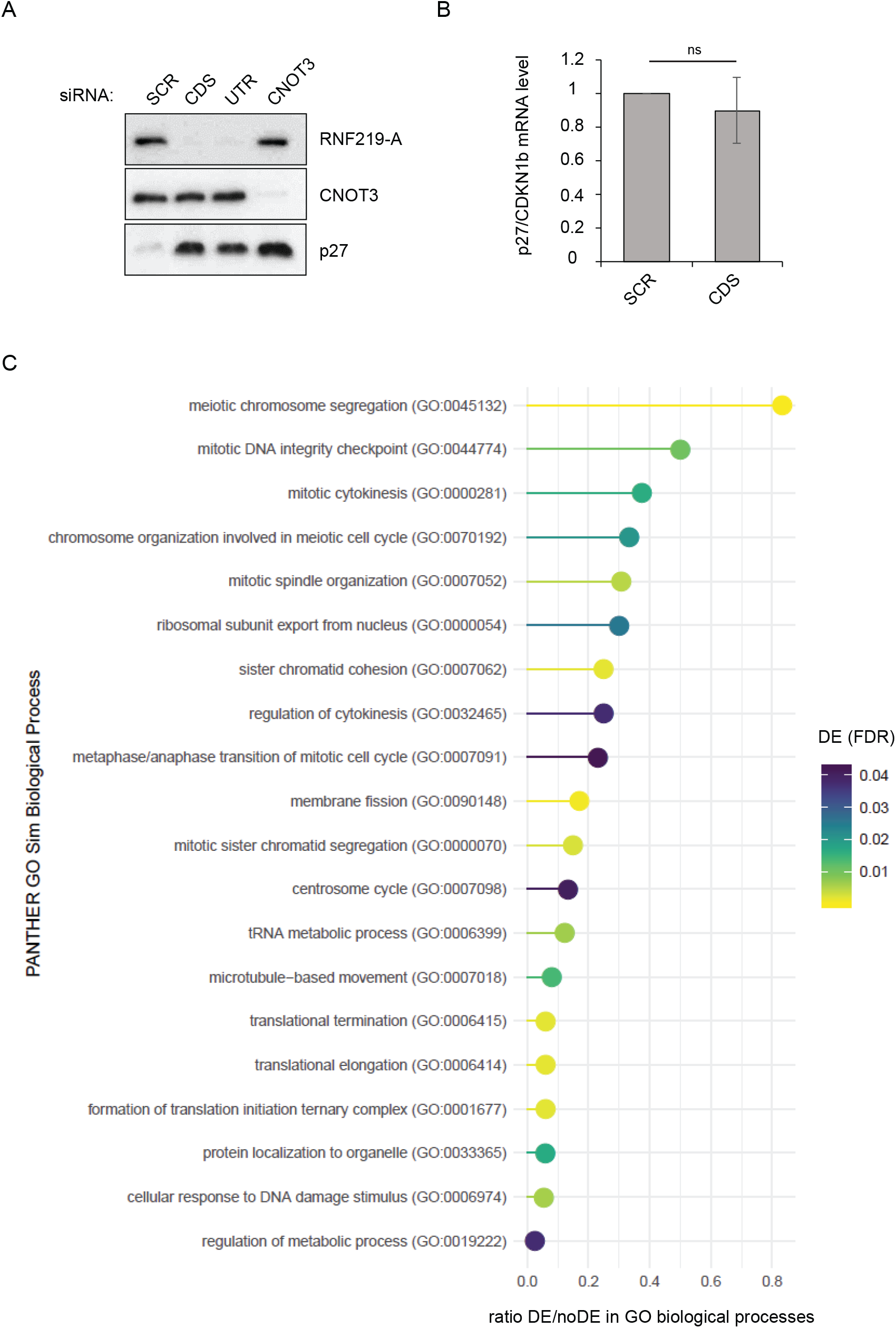
Endogenous RNF219 is implicated in cell cycle regulation. (A) Knock-down of RNF219 induces an increase in p27 protein level. HeLa cells were transfected with two siRNA against RNF219 (CDS and UTR) or CNOT3. RNF219, CNOT3 and p27 protein levels were analyzed by immunoblotting using RNF219-A, CNOT3 and p27 antibodies respectively. (B) Knock-down of RNF219 does not affect p27 mRNA level. RNA was extracted in control (SCR) or in RNF219 depleted cells (CDS). RT-QPCR was performed using primers specific to p27/CDKN1b (Table S1) and was normalized on 18S mRNA level. p27/CDKN1b mRNA level was set to 1 in the control (SCR). Error bars represent SD, n = 3. ns P > 0.05 based on unpaired two-tailed Student’s t-test. (C) GO term analysis of down-regulated genes in RNF219 depleted cells, using PANTHER Overrepresentation Test (which can be found on the geneontology.org website). The results of the GO term analysis are compiled in Table S5. DE stands for differentially expressed for the genes found down-regulated in RNF219 depleted cells (Table S3 and Table S4). noDE correspond to the genes of reference in the *Homo sapiens* genome used in the PANTHER test.

Next, we performed RNA sequencing in RNF219 depleted cells using two different siRNA (CDS and UTR2). The result of the differential expression analysis done between control (SCR) and RNF219 depleted cells (CDS or UTR2) show that 288 genes were found to be down-regulated genes while 80 were found upregulated (in the two siRNA used against RNF219; intersect of Table S3 and Table S4). GO term analysis of the down-regulated genes showed that RNF219 is generally implicated in cell cycle regulation and regulation of chromosome segregation (Fig. 6C, Table S5), with the most enriched GO term being meiotic chromosome segregation, mitotic DNA integrity checkpoint, mitotic cytokinesis, chromosome organization in meiotic cell cycle or mitotic spindle organization for instance.

Moreover, the GO term analysis also revealed that RNF219 is involved in translation initiation, elongation and termination, which correlates with our data showing that RNF219 affects mRNA expression by inhibiting translation in a yet unknown mechanism.

In conclusion, our results altogether suggest that RNF219 is implicated in cell cycle regulation. Moreover, its association with CCR4-NOT represses translation concomitantly to inhibition of deadenylation. We propose that RNF219 could serve as a molecular switch regulating pathway choices between different modes of translation repression (Fig. 7).

**Figure 7.**
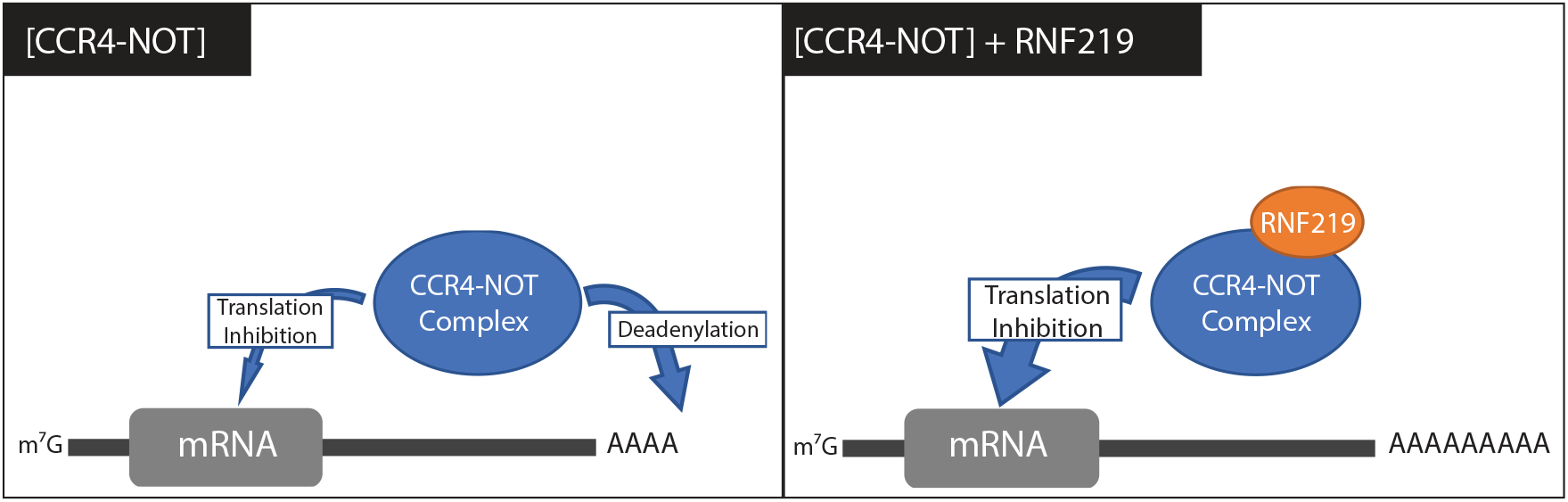
Schematic representation of RNF219 function. Recruitment of RNF219/CCR4-NOT complexes leads to mRNA translational repression in the absence of poly(A) tail deadenylation.

## DISCUSSION

In this study, we describe RNF219, a poorly characterized E3 ubiquitin ligase associated with the CCR4-NOT complex. Our biochemical data suggest that RNF219 stably associates with the CCR4-NOT complex (Fig. 2). The results indicate that RNF219 is not bound to the CCR4-NOT complex through the NOT module components CNOT2 or CNOT3, but either through direct or indirect binding to CNOT1 or other CNOT subunits.

The CCR4-NOT complex is a well-described key regulator of eukaryotic gene expression ^52^. Because numerous CCR4-NOT interacting partners regulate the activity of the complex and affect gene expression, RNF219 association with CCR4-NOT (Fig. 2) is an indication that it may also participate in these processes. Consistent with this hypothesis, we find that RNF219 inhibits targeted mRNA expression (Fig. 3) in a manner that greatly depends on its interaction with the CCR4-NOT complex. An exclusively cytoplasmic version of RNF219 is also able to repress targeted mRNA although to a lesser extent compared to WT RNF219 (Fig. 3E). This suggests that RNF219 has a yet unknown nuclear function in reducing mRNA expression. It is likely that it contributes to already described nuclear functions of CCR4-NOT such as transcription initiation, elongation, termination, splicing or export to the cytoplasm ^52^.

Strikingly, we found that RNF219 affects poly(A) tail shortening. This effect depends on RNF219 interaction with the CCR4-NOT complex (Fig. 5). Knowing that the predominantly described enzymatic activity of the CCR4-NOT complex is mRNA tail deadenylation, it is tempting to suggest that RNF219 acts as a negative regulator of the deadenylase activity of the complex. In their very recent study, Meijer et al. report that eiF4A2 and DDX6, two helicases compete to interact with CNOT1 ^53^. While DDX6 stimulates CNOT7 deadenylase activity, eiF4A2 is inhibitory in vitro. Interestingly longer poly(A) tails are bound to eiF4A2, similarly to poly(A) tails observed on the RNF219 targeted reporter ^53^. This data supports our model proposing specific interacting partners are able to switch CCR4-NOT function between different modes of translation repression, dependent or independent of deadenylation.

Recently, two studies reported that post-translational modifications of CCR4-NOT subunits affected its deadenylation activity ^47,54^. The first study, Cano et al. identified a E3 ligase, MEX-3C that interacts with the CCR4-NOT complex and promotes its deadenylase activity by ubiquitylation of the CNOT7 deadenylase ^47^. In the second study, Sharma et al. showed that deacetylation of CNOT7 by HDAC1 and 2 stimulates the deadenylase activity of the complex ^54^. Thus, post-translational modifications of the CCR4-NOT complex, including acetylation and ubiquitylation, modulate its activity. In contrast to previous reports, RNF219 is the first CCR4-NOT associated E3 ligase potentially inhibiting its enzymatic function. However, in our experiment, RNF219 ubiquitin ligase activity does not appear to be necessary for RNF219 mediated repression in the reporter assay. It is possible that the ubiquitin ligase activity serves at steps, such as RNF219 recruitment or residence time, absent in our tethering assay.

We showed that targeting RNF219 to a reporter mRNA on one hand inhibits its expression (Fig. 3) but on the other hand increased its poly(A) tail length (Fig. 5). This might seem surprising, as a long polyA tail has been classically associated with increased translation. However, recently Lima et al. demonstrated that in somatic cells highly expressed mRNAs tend to have short polyA tails ^8^. Consistently, our study reinforces the idea that there is no definite poly(A) tail length correlating with translation efficiency in somatic cells^7^.

In conclusion, in this study, we show that RNF219 is a novel factor implicated in post-transcriptional regulation of mRNA. Although, RNF219 mRNA targets are yet to be discovered, we found that RNF219 affects p27/CDKN1b expression post-transcriptionally, involving RNF219 in cell cycle regulation (Fig. 6).

To our knowledge RNF219 and eiF4A2 are the first identified modulators of CCR4-NOT activity preventing deadenylation and repressing translation at the same time ^53^. Interestingly, Youn et al. have recently described RNF219 and CCR4-NOT interaction using an in vivo proximity-dependent biotinylation (BioID) analysis ^55^. This report, defines the core components of stress granules (SGs) and P-Bodies (PBs). It is tempting to speculate that RNF219 modulates CCR4-NOT mediated deadenylation and translational repression in SG or PB to allow a rapid cellular response, enabling cells to start protein synthesis from already accumulated mRNAs. Future work will reveal physiologically relevant targets of the RNF219/CCR4-NOT complex and further characterize the pathway of translational repression it regulates.

## FUNDING

This work was supported by “FRM Amorçage” grant (AJE201229) to BS and FV, by a fellowship from ARC (PDF20130606877) to AG and by UMR9002 CNRS-University of Montpellier, ANR (ANR-15-CE12-0019-01 and ANR-17-CE12-0011-01), and AFM Telethon (17110) to MS.

## ACKNOWLEDGEMENT

We thank R. Pillai and W. Filipowicz for providing the expression plasmids of the tethering assay. P. Coulombe generously shared reagents and data prior to publication. P. Balaguer and A. Boulahtouf kindly provided the Firefly control vector and technical assistance. We acknowledge F. Bachand and A. David for protocols and help with polyribosome experiments. We thank members of the Molecular Virology laboratory for critically discussing data, sharing reagents, advice and support.

## Author Contributions

A.G., F.V. and B.S. conceived the study and analysed the data, with critical input from all the authors. A.G., F.V., A.C. and B.S. conducted experiments. A.G. and B.S. wrote the paper and all authors discussed the results and commented on the manuscript.

## Additional Information

### Supplementary information

includes four figures and two tables.

### Competing interest

The authors declare no competing interests.

