## Supplemental Material for "RNF219 regulates CCR4-NOT function in mRNA translation and deadenylation"

**Supplemental Figure S1**

**Supplemental Figure S2**

**Supplemental Figure S3**

**Supplemental Figure S4**

**Supplemental Table S1**

**Supplemental Table S2**

**Supplemental Table S3**

**Supplemental Table S4**

**Supplemental Table S5**

**Figure S1. RNF219 is a RING dependent ubiquitin ligase**

(A) The subcellular localization of RNF219 is preferentially either nuclear or cytoplasmic. Immuno-fluorescence performed with anti-HA on U2OS cells transfected with the FLAG-HA epitope-tagged RNF219 (FHA-RNF219) construct and nuclei were stained with Dapi (Scale bar, 12 μm). Statistical analysis of immuno-fluorescence experiment above. Error bars represent SD, n = 100.

(B) Immunoblot of extracts from HeLa cells transfected with control siRNA (SCR) or siRNA targeting the coding sequence (CDS) or the 3'UTR (UTR) of the RNF219 mRNA and probed with antibody against endogenous RNF219-A. The control condition (SCR) shows a band at 85 KDa, which is specifically depleted upon CDS and UTR siRNA transfection.

(C) Immunoblot of cell extracts from HeLa cells transfected with a CrispR/Cas9 plasmid expressing Cas9 and 2 selected guide RNA against RNF219 (CRISPR-1 and CRISPR-2). The pools represent mixed populations of transfected cells before clonal selection and Clones represent tested clones from single colony isolations derived from single cells.

(D) RNF219 was immuno-precipitated (IP) from HEK293T CRISPR-1 cells transfected with either pcDNA, a mock plasmid (mock) or FLAG-tagged RNF219 (WT) using the homemade RNF219-B antibody. The first 2 lanes contain the purification performed with an irrelevant IgG antibody. The presence of RNF219 in the IP was analyzed by immunoblotting using the RNF219-A and CNOT2 antibodies. The arrow indicates the band corresponding to CNOT2 while the asterisk indicates a non-specific band.

**Figure S2: RNF219 binds the CCR4-NOT complex**

(A) Endogenous CNOT3 (left), endogenous RNF219 (middle) or endogenous CNOT1 (right) was immuno-purified from HeLa cell extracts using an anti-CNOT3, anti-RNF219 (RNF219-B), anti-CNOT1 or irrelevant IgG antibody. IP were analysed by IB with indicated antibodies.

(B) RNF219 was immunoprecipitated using two different RNF219 specific antibodies (RNF219-B and RNF219-C) from HEK293T cell extracts. IP were analysed by IB with indicated antibodies. CCR4-NOT subunits were specifically enriched with both antibodies used for IP. IgG-B-PI (Pre-immune IgG antibody) and IgG-SC (a commercial control antibody) were used as control antibodies. The arrow indicates the band corresponding to CNOT2 while the asterisk indicates a non-specific band.

(C) RNF219 protein was truncated in multiple fragments until loss of CCR4-NOT interaction. RNF219 was truncated in six large fragments with a truncation step of 121 amino acids (represented by F1 to F6). The Flag-HA tagged fragments were transfected in cells. The F5 truncation lost interaction with three CCR4-NOT subunits (CNOT1, CNOT2, CNOT3). The Mock sample corresponds to cells transfected with a mock plasmid: pcDNA. IP were performed on FLAG beads from cell extracts and analysed by IB with indicated antibodies.

(D) The F5 truncation, which lost interaction with 3 CCR4-NOT subunits (CNOT1, CNOT2, CNOT3), was truncated in four fragments with a truncation step of 10 amino acids (represented by F5F, F5G, F5H, F5I). The Flag-HA tagged fragments were transfected in cells. The Mock sample corresponds to cells transfected with a mock plasmid: pcDNA. IP were performed on FLAG beads from cell extracts and analysed by IB with indicated antibodies.

(E) The interaction between RNF219 and the CCR4-NOT complex is RNase resistant. Top: the experimental scheme is shown. Bottom: FLAG and HA tagged RNF219 (FHA-RNF219) or FHA-Larp7 were IP'd on FLAG beads which were subjected to mock or RNase treatment. Co-IP'd proteins were analysed by IB. The RNase sensitive Larp7/CDK9 interaction served as positive control for RNase digestion.

**Figure S3. RNF219 affects the polyA tail length of a targeted mRNA**

(A) Immunoblot of experiment shown in Figure 5B.

(B) Renilla luciferase activity normalized to Firefly luciferase activity of experiment shown in Figure 5B.

**Figure S4. RNF219 affects the translation of a targeted mRNA**

(A) The inhibitory effect of RNF219 is proteasome independent. RL activity was measured in the indicated conditions. Renilla Luciferase (RL), Firefly (FL) plasmids and indicated NHA-protein constructs were co-transfected in HEK293T cells. The RL activity was measured and normalized for FL activity.

(B) Immuno-localization of NHA-RNF219 and NHA-RNF219-Cd. Immuno-fluorescence was performed with anti-HA antibody on HEK293T cells transfected with the indicated constructs and showed exclusively cytoplasmic staining for NHA-RNF219-Cd protein.

(C) Fractions of sucrose gradient are immunoblotted with anti-RNF219-A and anti-RPL3. Mono and polysomal fractions are indicated. Fractions 10-11 are used to measure the reporter mRNA level in the polysomal fraction (Poly) in Fig. 3F.

(D) Targeting RNF219 to the RL mRNA prevents recruitment of polyribosomes.

Polyribosome purification was done using the first protocol described in the methods. RL mRNA level in the Input and polyribosomal fraction (Poly) of extracts prepared from Hela cells transfected with NHA-LacZ or NHA-RNF219 normalized to GAPDH mRNA levels.

(E) Translation efficiency of Hela cells is monitored by quantifying Puromycin incorporation in nascent protein. Immunoblot using anti-puromycin antibody is performed on total cell extract after puromycin pulse (Goodman and Hornberger, 2013). In the mock condition, no puromycin is added to the medium. In the second lane, the translation inhibitor Cycloheximide is added at the same time for 30 min. In the third and fourth lanes, puromycin pulses were of 15 and 30-min respectively.

**Supplementary Table S1:**

List of primers used in this study

**Supplementary Table S2:**

List of RNF219 interacting proteins detected by MS/MS with percent peptide coverage by amino acid count above 22%. CCR4-NOT subunits are highlighted in yellow. Known CCR4-NOT recruiting proteins are highlighted in blue.

**Supplementary Table S3:**

Results of the differential expression analysis between control (SCR) and RNF219 depleted cells (CDS). The differential expression analysis was done using Cufflinks (http://cufflinks.cbcb.umd.edu/) at the RNA sequencing platform Fasteris SA.

**Supplementary Table S4:**

Results of the differential expression analysis between control (SCR) and RNF219 depleted cells (UTR2). The differential expression analysis was done using Cufflinks (http://cufflinks.cbcb.umd.edu/) at the RNA sequencing platform Fasteris SA.

**Supplementary Table S5:**

Results of the GO term analysis done using the list of down-regulated genes in RNF219 depleted (combination of down-regulated genes found in common in siCDS and siUTR2 treated cells), using PANTHER Overrepresentation Test (which can be found on the geneontology.org website).

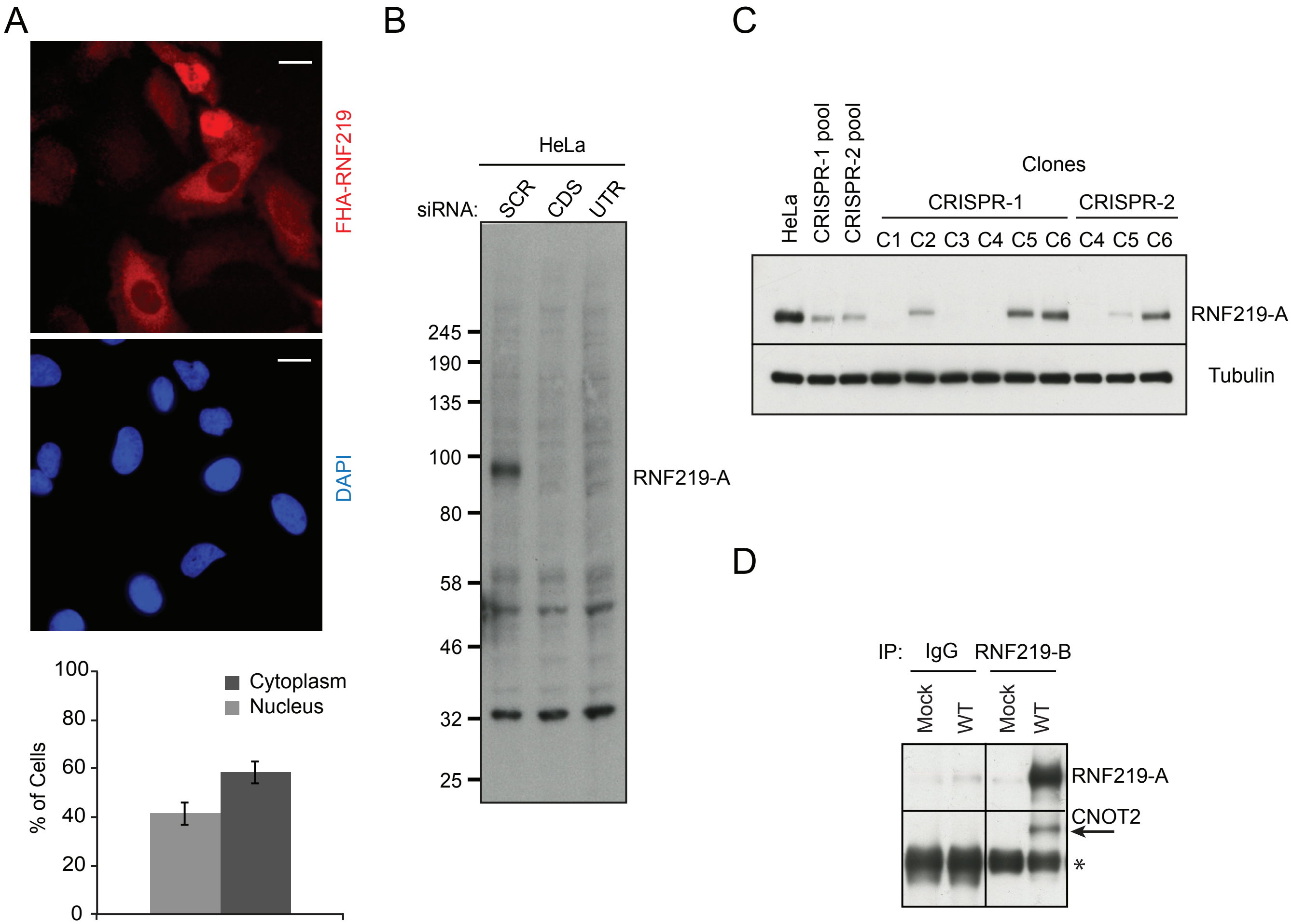
A
FHA-RNF219
DAPI
Cytoplasm
Nucleus
% of Cells
100
80
60
40
20
0
B
HeLa
siRNA:
SCR
CDS
UTR
245
190
135
100
80
58
46
32
25
RNF219-A
C
HeLa
CRISPR-1 pool
CRISPR-2 pool
Clones
CRISPR-1
CRISPR-2
C1 C2 C3 C4 C5 C6 C4 C5 C6
RNF219-A
Tubulin
D
IP:
IgG
RNF219-B
Mock
WT
Mock
WT
RNF219-A
CNOT2
*

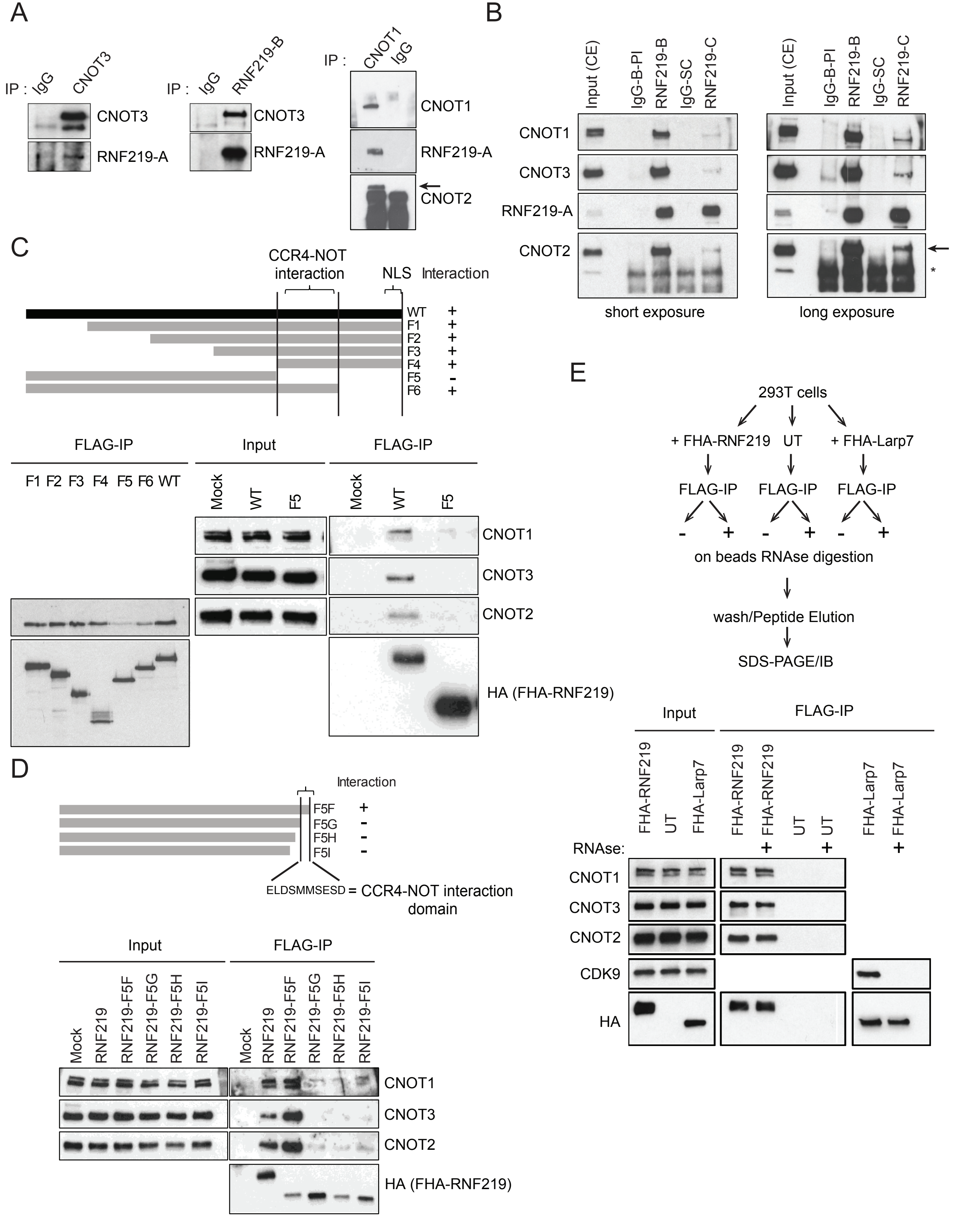

A
IP : IgG CNOT3
CNOT3
RNF219-A
IP : IgG RNF219-B
CNOT3
RNF219-A
IP : CNOT1 IgG
CNOT1
RNF219-A
CNOT2
B
Input (CE) IgG-B-PI RNF219-B IgG-SC RNF219-C
CNOT1
CNOT3
RNF219-A
CNOT2
short exposure
long exposure
*
C
CCR4-NOT interaction
NLS
Interaction
WT +
F1 +
F2 +
F3 +
F4 +
F5 -
F6 +
FLAG-IP
F1 F2 F3 F4 F5 F6 WT
Input
Mock WT F5
FLAG-IP
Mock WT F5
CNOT1
CNOT3
CNOT2
HA (FHA-RNF219)
D
Interaction
F5F +
F5G -
F5H -
F5I -
ELDSMMSESD = CCR4-NOT interaction domain
Input
Mock RNF219 RNF219-F5F RNF219-F5G RNF219-F5H RNF219-F5I
FLAG-IP
Mock RNF219 RNF219-F5F RNF219-F5G RNF219-F5H RNF219-F5I
CNOT1
CNOT3
CNOT2
HA (FHA-RNF219)
E
293T cells
+ FHA-RNF219 UT + FHA-Larp7
FLAG-IP FLAG-IP FLAG-IP
- + - + - +
on beads RNAse digestion
wash/Peptide Elution
SDS-PAGE/IB
Input
FLAG-IP
FHA-RNF219 UT FHA-Larp7 FHA-RNF219 FHA-RNF219 UT UT FHA-Larp7 FHA-Larp7
RNAse: + + +
CNOT1
CNOT3
CNOT2
CDK9
HA

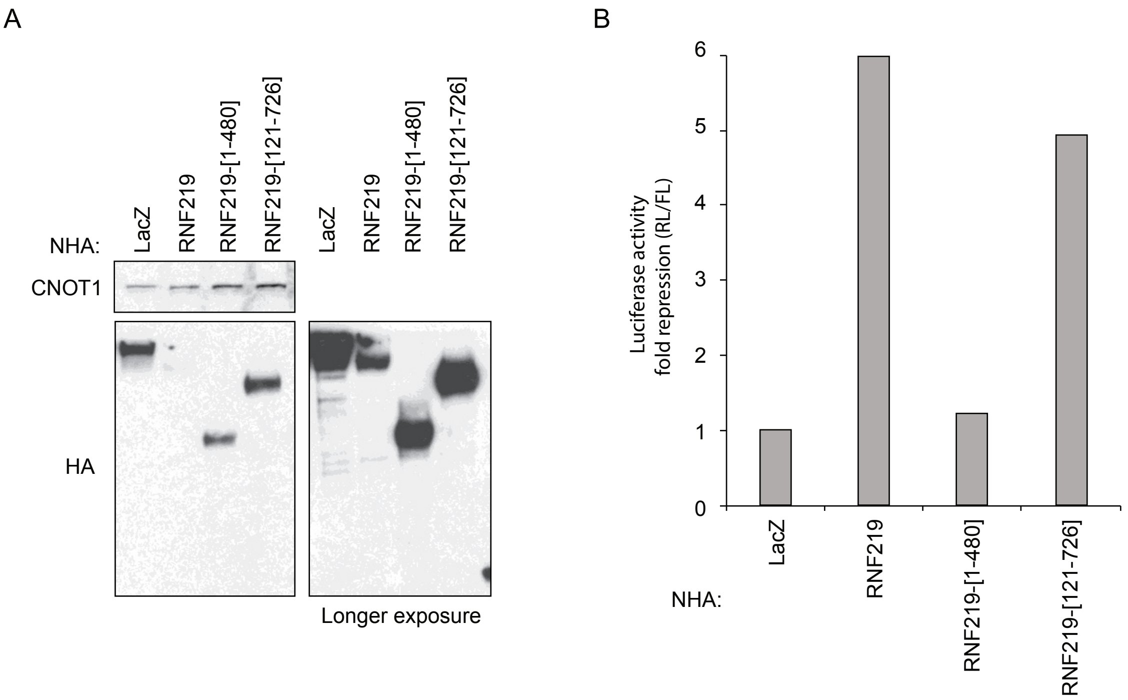

A
NHA:
LacZ
RNF219
RNF219-[1-480]
RNF219-[121-726]
LacZ
RNF219
RNF219-[1-480]
RNF219-[121-726]
CNOT1
HA
Longer exposure
B
Luciferase activity
fold repression (RL/FL)
0
1
2
3
4
5
6
NHA:
LacZ
RNF219
RNF219-[1-480]
RNF219-[121-726]

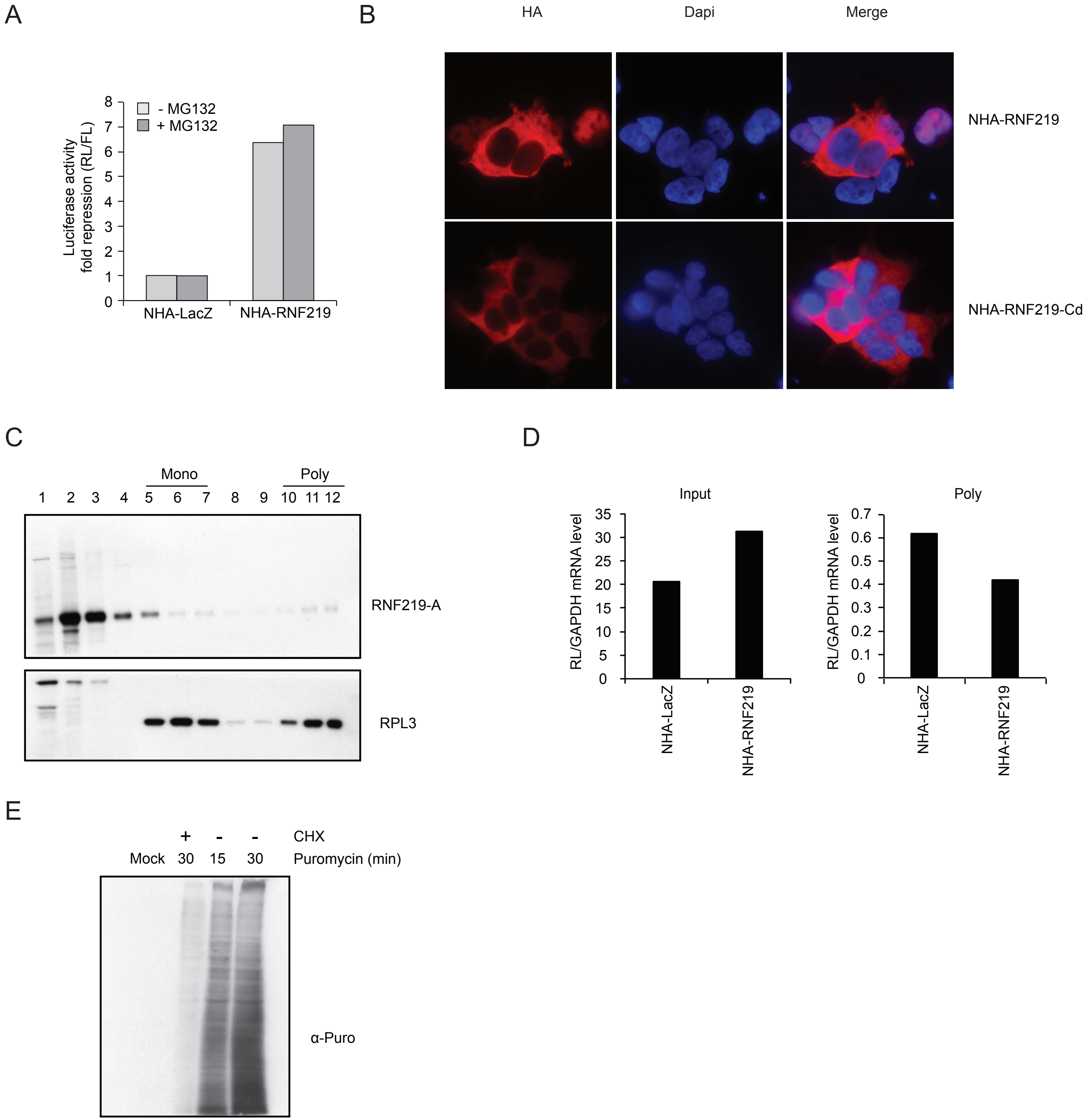
A
B
HA
Dapi
Merge
- MG132
+ MG132
Luciferase activity
fold repression (RL/FL)
8
7
6
5
4
3
2
1
0
NHA-LacZ
NHA-RNF219
NHA-RNF219
NHA-RNF219-Cd
C
Mono
Poly
1 2 3 4 5 6 7 8 9 10 11 12
RNF219-A
RPL3
D
Input
Poly
RL/GAPDH mRNA level
35
30
25
20
15
10
5
0
0.7
0.6
0.5
0.4
0.3
0.2
0.1
0
NHA-LacZ
NHA-RNF219
E
+ - - CHX
Mock 30 15 30 Puromycin (min)
α-Puro

Supplemental Table S1

| | | |
|---|---|---|
| RTqPCR | RNF219 F | CATCACGTGCCACATTTGCT |
| | RNF219 R | ATGCTTCCTGACCGTATGGC |
| | GAPDH F | ACAGTCCATGCCATCACTGCC |
| | GAPDH R | GCCTGCTTCACCACCTTCTTG |
| | RENILLA F | TCGTCCATGCTGAGAGTGTC |
| | RENILLA R | CTAACCTCGCCCTTCTCCTT |
| | FIREFLY F | TCAAAGAGGCGAACTGTGTG |
| | FIREFLY R | TTTTCCGTCATCGTCTTTCC |
| | p27/CDKN1b F | GGGGAAGGGTTTGAATTGTT |
| | p27/CDKN1b R | GGAGGGCAGTGAGGATAGGT |

| | | |
|---|---|---|
| SiRNA | SCRAMBLE | UCUGCAAGGUUAGGCGUCU |
| | CNOT1_A | GGAACUUGUUUGAAGAAUA |
| | CNOT1_B | CAAGGUCCUUGGUAUAGUA |
| | RNF219 CDS | GCAGACCUUAACUGUUCUA |
| | RNF219 UTR | CUUGUCUUCAGAACUUGGA |
| | RNF219 UTR2 | CTTGTCTTCAGAACTTGGA |

| | | |
|---|---|---|
| ePAT | LUC_2 | GCTTTATTTGTGAAATTTGTGATGC |
| | LUC_3 | CGAGCAGACATGATAAGATACATTG |
| | GAPDH | GAGCCGCACCTTGTCATGTA |

Supplemental Table S2

| UniProt | Gene | Amino acids | % Peptide coverage |
|---|---|---|---|
| Q9UFF9 | CNOT8 | 292 | 63 |
| P62258 | 1433E | 255 | 58 |
| Q9C0C2 | TB182 | 1729 | 56,9 |
| Q5W0B1 | RN219 | 726 | 55,9 |
| Q9NZN8 | CNOT2 | 540 | 55,7 |
| Q92600 | CNOT9 | 299 | 50,8 |
| Q8IY67 | RAVR1 | 606 | 49,1 |
| A5YKK6 | CNOT1 | 2376 | 48,4 |
| P40937 | RFC5 | 340 | 47,9 |
| Q9UIV1 | CNOT7 | 285 | 46,3 |
| P33993 | MCM7 | 719 | 45,2 |
| P52272 | HNRPM | 730 | 44,8 |
| Q14247 | SRC8 | 550 | 42,2 |
| Q9UKZ1 | CB029 | 510 | 41 |
| Q9H9A5 | CNOT10 | 744 | 40,2 |
| P35250 | RFC2 | 354 | 39,5 |
| Q14106 | TOB2 | 344 | 36,3 |
| Q9ULX6 | AKP8L | 646 | 36,2 |
| P02545 | LMNA | 664 | 35,4 |
| Q8IXQ3 | CI040 | 194 | 35,1 |
| Q9NWU2 | CT011 | 228 | 34,6 |
| Q86VI3 | IQGA3 | 1631 | 34,3 |
| Q14134 | TRI29 | 588 | 34,2 |
| P49368 | TCPG | 545 | 34 |
| Q9ULM6 | CNOT6 | 557 | 33 |
| P10398 | ARAF | 609 | 31,5 |
| Q9HC44 | GPBL1 | 474 | 31,2 |
| Q9HAU0 | PKHA5 | 1116 | 30,6 |
| Q96LI5 | CNOT6L | 555 | 29,7 |
| Q8WUF5 | IASPP | 828 | 29 |
| Q6P1J9 | CDC73 | 531 | 28,8 |
| P62333 | PRS10 | 389 | 28,8 |
| Q14201 | BTG3 | 252 | 28,2 |
| P50616 | TOB1 | 345 | 28,1 |
| Q9NXR1 | NDE1 | 346 | 28,1 |
| P27708 | PYR1 | 2225 | 28 |
| O00233 | PSMD9 | 223 | 27,4 |
| Q92974 | ARHG2 | 986 | 26,5 |
| Q9BSD7 | CA057 | 190 | 26,3 |
| P40938 | RFC3 | 356 | 26,1 |
| P62195 | PRS8 | 406 | 26,1 |
| P40227 | TCPZ | 531 | 26 |
| P35998 | PRS7 | 433 | 25,9 |
| Q14244 | MAP7 | 749 | 25,8 |
| Q15208 | STK38 | 465 | 25,6 |
| P50750 | CDK9 | 372 | 25,3 |
| P04049 | RAF1 | 648 | 24,7 |
| O00571 | DDX3X | 662 | 24,5 |
| O75175-1 | CNOT3 | 753 | 22,4 |
| Q9ULX6 | AKP8L | 646 | 22,4 |
